# Modes of interaction between individuals dominate the topologies of real world networks

**DOI:** 10.1101/011288

**Authors:** Insuk Lee, Eiru Kim, Edward M. Marcotte

## Abstract

We find that the topologies of real world networks, such as those formed within human societies, by the Internet, or among cellular proteins, are dominated by the mode of the interactions considered among the individuals. Consequently, a major dichotomy in previously studied networks arises from modeling networks in terms of pairwise versus group tasks. The former often intrinsically give rise to scale-free, disassortative, hierarchical networks, whereas the latter often give rise to broad-scale, assortative, nonhierarchical networks. These dependencies explain contrasting observations among previous topological analyses of real world complex systems. We also observe this trend in systems with natural hierarchies, in which alternate representations of the same networks, but which capture different levels of the hierarchy, manifest these signature topological differences. For example, in both the Internet and cellular proteomes, networks of lower-level system components (routers within domains or proteins within biological processes) are assortative and nonhierarchical, whereas networks of upper-level system components (internet domains or biological processes) are disassortative and hierarchical. Our results demonstrate that network topologies of complex systems must be interpreted in light of their hierarchical natures and interaction types.

## Introduction

Networks occur in many contexts in the real world, and the topologies of these real-world networks have potentially large practical impact in areas of biological, social, and technical importance. Topological implications of human social networks, for example, influence public policies (such as for the development of effective vaccination schemes[1]) and business strategies (such as for allocating marketing resources[2]). Likewise, the topology of the Internet affects routing protocols for robust and cost-effective communications[3], and in biology, the topologies of protein-protein interaction (PPI) networks have informed our understanding of cells and organisms[4].

As a consequence, network topologies have been extensively characterized with respect to their global topological properties, such as node degree distributions[5], node hierarchical organization[6], and assortativity (the degree correlation between connected nodes)[7]. Differences in such properties have been noted for many real world networks[8]. In the course of studying networks, we realized that many of these historical observations of contrasting network topologies could be explained by a simplifying model: that most real world networks can be categorized as one of two major classes of networks – those capturing intrinsically pairwise activities (e.g., dating or pairwise physical interactions between proteins) and those capturing intrinsically group activities (e.g., boards of directors of companies or membership in the same protein complexes). In this paper, we demonstrate that this distinction explains many of the major topology differences amongst social networks, the Internet, and biological networks, and that networks generated by the same class of activities – regardless of the precise nature of those activities – often have similar topological properties.

## Results and Discussion

### An intrinsic dichotomy between contact- and task-centric networks

We illustrate this key distinction among the two network classes in Figure 1 by introducing toy examples of two types of human social interactions similarly composed of 11 people (nodes) organized into three groups (indicated by node colors). Interpersonal relationships (edges) might be based on direct personal contact—the *contact-centric model* (Figure 1a)—such as for the cases of online dating[9] or sexual contacts[10], or alternatively based on sharing roles to perform a common task—the *task-centric model* (Figure 1b)—such as for sharing membership on company boards[11] or co-authorship of scientific papers[12]. In the contact-centric network, a few attractive individuals (represented as squares) have a large number of partners (Figure 1c). In contrast, a task-centric network is characterized by group activities in which the pairwise interaction reflects the tendency for individuals to participate in the same groups. Note that individuals may participate in the same task but never actually directly contact each other (represented as dotted lines)—e.g., many coauthors for scientific papers do not have a personal relationship (Figure 1d). The networks’ topologies can be characterized by various global topological properties[8]. Here, we will consider the three most widely studied topological properties: node degree distribution, node hierarchical organization, and assortativity.

**Figure 1.**
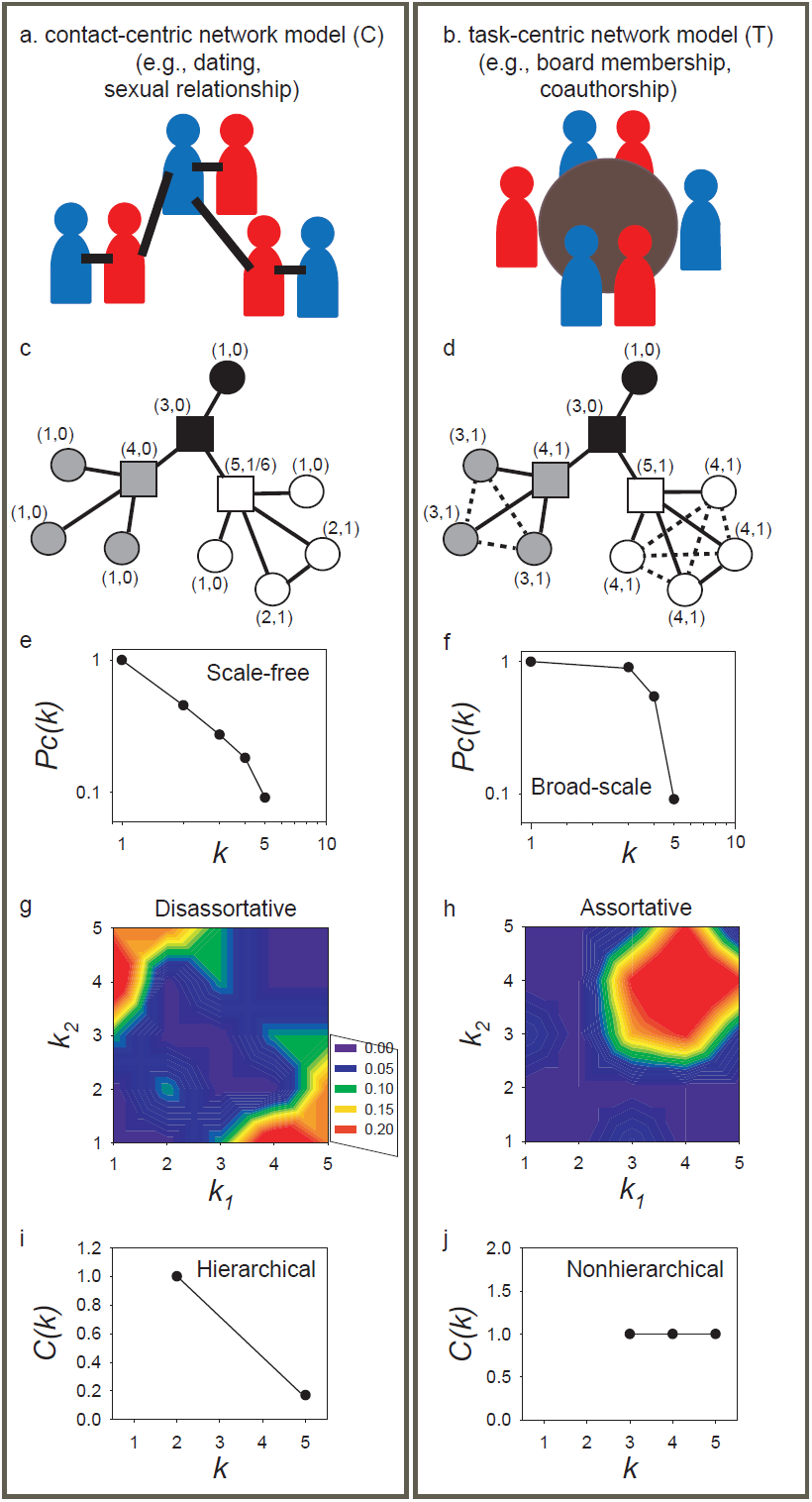
Two broad classes of networks naturally occur in real world settings based on whether interactions between entities are intrinsically pairwise (left panel) or based on groups (right panel). Examples in human social networks might include (a) dating or sexual relationships, and (b) board memberships or co-authorships of scientific papers. (c and d) indicate small toy examples for these two types of human social networks, each composed of 11 individuals (nodes) organized into three groups (indicated by node colors). Edges in (c) indicate direct contacts between persons; edges in (d) indicate participation in shared tasks. Numbers in (c) denote the degree connectivity (count of associated edges) and clustering coefficient[15] for each node. Individuals with high node degree are marked as squares. In spite of their simplicity, the two toy networks show distinct topological features. Their cumulative probability distributions of nodes with >*k* degree (*Pc*(*k*)) differ; (e) is scale-free, (f) is broad-scale. They differ in being (g) disassortative or (h) assortative networks, as seen by heat-map representations of the enrichment for connections between nodes of varying degrees. Finally, they exhibit (i) hierarchical or (j) nonhierarchical network topologies, as judged by their relationships between node degree connectivity (*k*) and node clustering coefficients (*C*(*k*)).

An examination of the topologies of these toy networks reveals striking differences. In general, the distribution of node degree of scale-free networks approximates a power-law for the entire range of node degrees[13], *Pc*(*k*) = *k*^-*γ*^, where *Pc*(*k*) represents the cumulative probability of having nodes with >*k* degree. The contact-centric network is expected to be more scale-free in nature, because there is no constraint limiting the number of links for individual nodes by this modeling perspective, permitting few nodes with many partners, while most nodes have only few partners (Figure 1e). In contrast, the task-centric network is expected to be of broad-scale[5] (Figure 1f). A broad-scale network is often characterized by a power-law regime followed by exponential trimming of degree, represented by the function *Pc*(*k*) = *k*^-*γ*^*exp*(-*k*/*k*_0_), where *k_0_* is the cutoff degree for exponential decay. In general, networks following this truncated power-law function exhibit an increased proportion of nodes with medium-connectivity, probably due to the constraint on the number of links by sizes of task-groups of the networks, resulting in reduced numbers of both hubs and nodes with low-connectivity. Because of the connections among entities performing the same tasks, the task-centric network shows a degree preference corresponding to the preferred sizes of the groupings. Thus, these two network classes intrinsically give rise to distinct node degree distributions.

We see an equally strong dichotomy for a second well-characterized topological property that of the correlation in degree between connected nodes. A given network is termed assortative if its hub-hub connections are enriched; if depleted, the network is termed disassortative[7]. The assortativity of networks can be measured by the statistical significance of the enrichment in connections between various ranges of node degrees as compared to permuted networks[14], and can be visualized as a heat map. Notably, the contact-centric network example is disassortative (Figure 1g), while the task-centric network example is assortative (Figure 1h).

The third major topological property that we consider is the hierarchical organization[6]. In a hierarchically organized network, hub components bridge many disconnected regions of the network to efficiently coordinate all system components. The number of these far-reaching connections decreases as the node degree decreases, entailing increasing proportions of connections toward local neighbors. This indicates a hierarchical contribution of system components with hub components generally sharing high betweenness (a measure of network centrality). As a consequence, the clustering coefficients *C(k)*[15] of nodes decrease as their node degree *k* increases in hierarchically organized networks. It is noticeable that this definition of hierarchy assumes that real world networks are in general scale-free and modular. Thus, disassortative networks are also expected to be hierarchical networks (i.e., assortativity and hierarchy are not independent topological features). There are other hierarchy models which are independent from network modularity[16,17]. However, the hierarchy by *C(k)* decreasing with *k* has been widely used for previous network topological studies we discussed in this paper. Many real world networks exhibit this signature of hierarchical organization[6], as does the contact-centric toy network example in Figure 1i. Nonhierarchical networks, however, exhibit roughly equal clustering coefficients, regardless of node degree. This trend suggests enriched connections between components with similar numbers of network neighbors, and the task-centric network example is accordingly nonhierarchical (Figure 1j).

Thus, these two major classes of networks, characterized by the nature of the relationships among the interacting entities, show distinct topological properties: a contact-centric network is a scale-free, hierarchical, disassortative network; a task-centric one is a broad-scale, nonhierarchical, assortative network. The analyses of these toy examples suggest that the contrasting topological properties of many real world networks might also stem from this intrinsic dichotomy of network type. We thus next examined real world networks to test this hypothesis (listed in Supplementary Table 1).

### Real-world networks exhibit the same dichotomy

We first compared the three global topological properties for two human social networks, analyzing a contact-centric online dating network[9] and a task-centric network of boards of directors of US companies[11] (Figure 2, **top panel**). The node degree distribution of the dating network showed more scale-free character, while that of the board of directors network showed broad-scale (Figure 2a). We observed contrasting assortativity between the two networks: the dating network is disassortative, but the board of directors network is assortative (Figure 2b). Similarly, we found a clearly hierarchical organization for the dating network but a nonhierarchical organization for the director board network (Figure 2c). Thus, as for the toy network examples in Figure 1, real world human social networks also show contrasting topologies according to the distinction between intrinsically pairwise vs. group activities. In the online dating network, most people online date only a few partners, while a few individuals make large numbers of online dating contacts. In contrast, membership on boards of directors carries significant obligations. Thus, directors typically participate in only a limited number of boards, and boards are limited in size; as a consequence, clustering coefficients are often similar between members belonging to large boards and those who belong to small ones.

**Figure 2.**
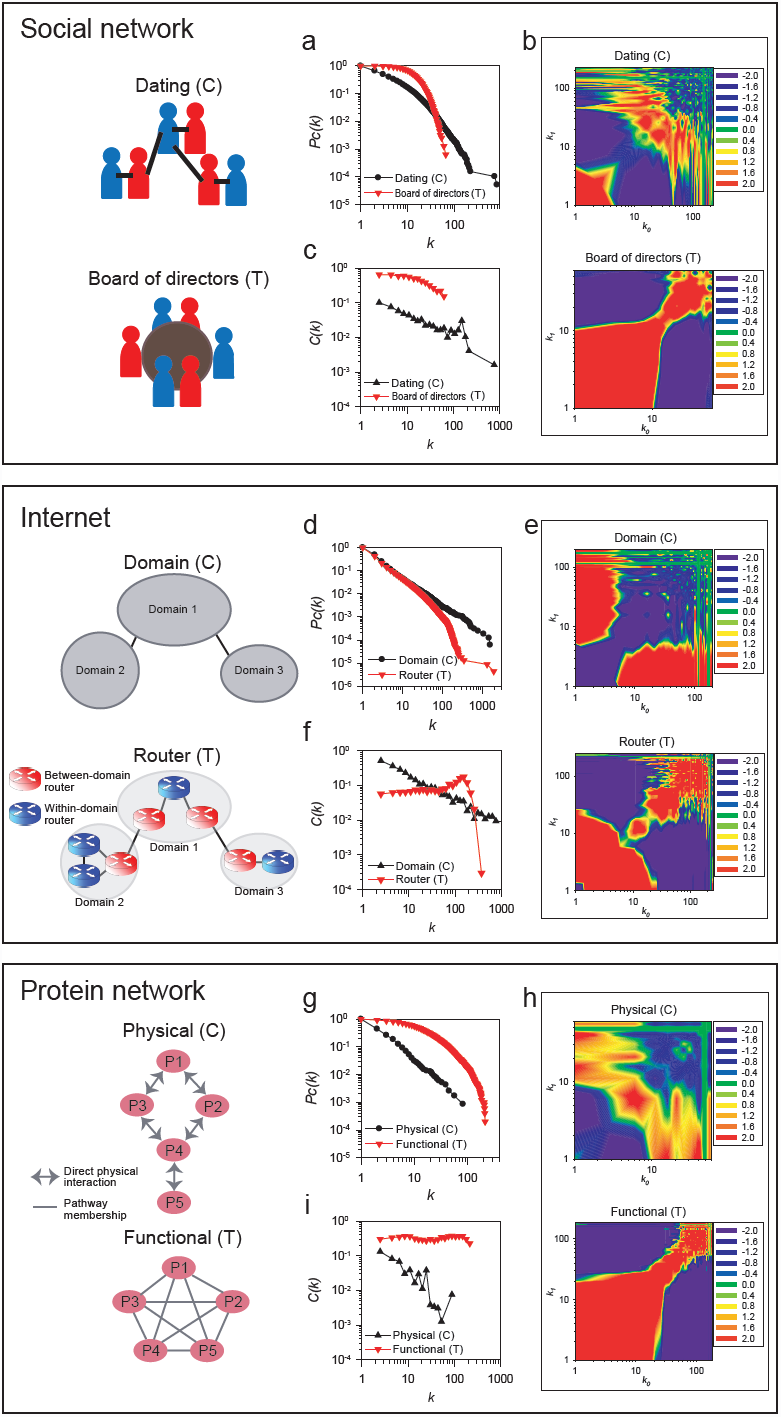
Topological analysis of three real world networks (detailed in Supplementary Table 1). We analyzed contact-centric networks (indicated by the letter C) and task-centric networks (indicated by the letter T) for their (a, d, g) degree distribution, (b, e, h) assortativity, and (c, f, i) hierarchical structure. We tested two different human social networks, a contact-centric online dating network (Dating)[9] and a task-centric network of shared membership on US company boards of directors (Director board)[11]. Similarly, we analyzed the Internet at two different levels, the router-level (Router) and domain level (Domain). The router-level Internet is task-centric and the domain-level Internet is contact-centric. Finally, we analyzed two alternate protein networks, testing pairwise protein interactions (CCSB-YI1)[20] and functional protein interactions (YeastNet core)[21], as examples of contact-centric and task-centric protein networks, respectively. We measured the cumulative probability of nodes with degree >*k* (*Pc*(*k*)) for the full range of node degree[13] to test scale-freeness of networks. We measured hierarchical connectivity by testing for decreasing clustering coefficients (*C*(*k*)) as a function of increasing node degree (*k*). Network assortativities were visualized as heat maps of the enrichment of connections between various ranges of degrees compared to permuted networks[14]; red indicates enrichment and blue indicates suppression of connectivity. In every case, real-world networks showed topological properties consistent with being either contact-or task-centric, as appropriate.

The Internet represents another widely studied real world complex system comprising computers and other devices with IP addresses that communicate *via* a network of routers, each implementing routing protocols. Internet scientists have considered two major networks, those composed of routers and those of domains (or autonomous systems) (Figure 2, middle panel). A domain is an entity containing multiple routers under control of a network operator(s) (e.g., Internet Service Provider; ISP) using a common routing policy to the Internet. Thus, each domain is considered to be an operational unit component of the Internet. For each domain, within-domain routing protocols and between-domain protocols such as the Border Gateway Protocol (BGP) can be implemented within one or more separate routers. Therefore, the Internet, when considered at the domain-level, maps only peer-to-peer connections between BGP routers, while the router-level maps all router connections including both within-domain and between-domain connections.

Thus, the domain-level Internet model can be considered to be a contact-centric network, while the router-level Internet model resembles the task-centric model. Accordingly, analysis of published router network and domain network[18] revealed the topological properties consistent with the contact-centric and task-centric models, just as for the human social networks. The contact-centric domain network is scale-free, disassortative, and hierarchical, but the task-centric router network is broad-scale, assortative, and nonhierarchical (Figure 2d-f). Disassortativity of a domain-level Internet has been also observed in a previous study[19]. Routers with more than 300 degree connections do show exceptionally low clustering coefficients (Figure 2f, highest degree nodes). However, this pattern only affects a small number of routers at the highest level domain, and the overall two levels of organization share topological characteristics consistent with the trends expected for task-centric models. This trend arises because the Internet has top-level domains for mediating communication between lower-level domains. Between-domain routers of the top-level domains generally have connections to a large number of lower-level domains, resulting in much lower clustering coefficients for these few hub routers than for the rest of routers in the Internet. Thus, the Internet, considered at the level of organization for the entire router network, is nonhierarchical.

Similarly, biologists have measured interactions among cellular proteins, describing complex biological networks. These networks naturally fall into contact or task centric models. For example, measurements employing the yeast two-hybrid (Y2H) methodology map the direct physical pairwise contacts between proteins. In contrast, proteins participate in groups—in pathways and complexes—in order to fulfill their functional roles within cells, and physical interactions are but one indication of functional association. We can accordingly model PPI networks considering either direct physical contacts (contact-centric) or functional associations (task-centric) between proteins (Figure 2, **bottom panel**). We analyzed a high-confidence genome-wide Y2H map of yeast proteins, CCSB-YI1[20] as a representative physical interaction network, and a published functional protein network (YeastNet core)[21] as a representative functional network. Consistent with the contact/task dichotomy, the physical network is scale-free[22], disassortative[14], and hierarchical[22], while the functional network is broad-scale, assortative, and nonhierarchical (Figure 2g-i). Thus, in protein networks, as in social and technology networks, the contact/task centric models explain the dominant topological properties.

These models can account for contrasting observations from previous topology studies for a variety of real world networks. Typically, many PPI networks have been implicitly modeled from the contact-centric view, while the task-centric view has been implicitly adopted for modeling many human social networks. Thus, PPI networks are generally thought to be scale-free disassortative hierarchical networks, and the human social networks as broad-scale assortative nonhierarchical networks. However, there have been reports of noncanonical topological properties for protein functional networks[21,23] and online dating networks[9]. These inconsistencies are easily explained as resulting from the contact/task centric dichotomy. Mixtures of these two models, for example by combining Y2H protein interactions and protein complexes into a single network model, may have led to inconsistent observations regarding the suppression of hub-hub connections in different PPI networks[14,24]. In fact, as we next show, the choice of how the same entities are represented in a network model markedly affects the network topology.

### Alternate network representations of the same entities can also exhibit this dichotomy

A study of the toy examples in Figure 1 immediately suggests that different representations of the associations among the same entities could give rise to networks with either topology. We see that this is indeed the case in real-world networks and can be easily illustrated for both the internet and protein networks. In both cases, the topological properties of the resulting alternate network models match those expected for contact and task centric networks.

For example, for the case of the internet, the router and domain networks differ in not only edge representation but also node representation—the task-centric router-level Internet has nodes of individual routers, whereas the contact-centric domain-level Internet has nodes of domains composed of multiple routers. As seen above (Figure 2d-f; reprinted in Figure 3a for clarity), the domain network is scale-free, disassortative, and hierarchical, but the router network is broad-scale, assortative, and nonhierarchical. This observation of contrasting Internet topology between router-and domain-level also has been reported by[18]. Likewise, cellular proteins can be modeled by grouping according to pairwise protein interactions or, alternatively, by clustering the proteins into functional modules or biological processes. We therefore defined biological processes by grouping proteins that were highly connected to one another in a functional network. From the resulting 333 biological processes defined by hierarchical clustering, we generated a network of processes as described in[25]. The resultant process network was revealed to be disassortative (Supplementary Figure 1) and hierarchical (Figure 3b). Regardless of node degree, most proteins and routers showed clustering coefficients near the average clustering coefficient for the entire protein or router network. Notably, our findings were consistent when biological process networks were created using alternate methods (e.g. using an alternate clustering algorithm, MCL[26], or using pre-existing process annotations from the Gene Ontology[27]) (Supplementary Figure 2). Thus, these observations thus suggest common rules-of-organization between the Internet and the yeast proteome at equivalent system levels—assortative and nonhierarchical organization among lower-level components, and disassortative and hierarchical organization among upper-level components (Figure 3a **and** 3b).

**Figure 3.**
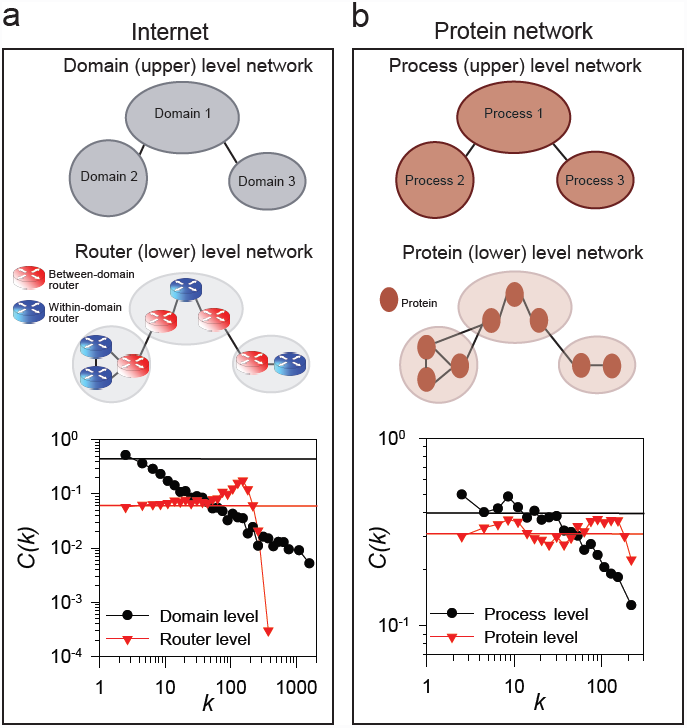
Alternate representations of the same network can lead to different topologies, especially for networks with natural hierarchical organization. We illustrate this tendency for (a) the Internet and (b) the yeast cell proteome. Each can be modeled by networks at two different granularities, representing nodes either as upper level components (internet domains or protein processes) or lower level components (internet routers or individual proteins). For the internet, previous Internet mapping studies provide both a router-level network and a domain-level network[18]; each domain is composed on multiple routers, and domains are connected *via* between-domain routers. For the protein network, we defined protein processes by hierarchically clustering proteins based on their pairwise interactions as in[25]. A total of 333 biological processes were defined and connections between processes were defined based on pairwise interactions between proteins within each process. The networks’ hierarchical structure was analyzed and plotted as in Figure 2, marking the mean clustering coefficient for each entire network as a horizontal solid line in the plot. The non-hierarchical router and protein networks generally exhibited clustering coefficients near this average regardless of node degree, although for the Internet router-level network, routers with >300 connections showed exceptionally low clustering coefficients primarily due to a small number of between-domain routers located at a few top-level domains of the Internet.

This intrinsic dichotomy between pairwise and group-based interactions helps to clarify many previous network analyses. It has been argued that the enrichment of hub-hub connections in the board of directors network may suggest the existence of a *super-group* of decision-makers that influences the business of the entire nation[11]. However, given that this network represents a task-centric model, we would expect this property to be an intrinsic feature of the model. In contrast to the board of directors network, hub-hub connections are suppressed in the contact-centric online dating network (and similarly in sexual contact networks[28]). This trend has been argued as an example of the robustness of social networks, since enhanced contacts between hubs would dramatically increase the risk to society, as for example, might result from hub-hub transmission of sexually transmitted diseases[10]. Similarly, physical protein interaction networks have been argued to exhibit robustness on the basis of their disassortative topology by localizing the effects of deleterious protein perturbations[14]. However, in both of these cases, a consideration of contact vs. task centric networks reveals that the rule of dissortativeness-robustness applies only to contact-centric networks, as task-centric networks are intrinsically assortative.

## Conclusions

In summary, we describe two broad classes of network models whose intrinsic topological properties explain observations on many real world networks, as we have shown with examples from human societies, the Internet, and cellular proteins. Contact-centric networks are characterized by intrinsically pairwise interactions, generating scale-free, disassortative, hierarchical networks. Task-centric networks are characterized by interactions within and between groups, and give rise to broad-scale, assortative, nonhierarchical networks. Alternative representations of the same entities, such as by grouping individuals together hierarchically, can also give rise to such networks, with lower-level representations generating assortative, nonhierarchical networks, and upper-level, clustered representations generating disassortative, hierarchical networks. The intrinsic topologies of these two classes of networks may account for previous paradoxical observations among real world networks, and imply similar organizational principles among human societies, the Internet, and cellular proteomes.

## Methods

Global topological analyses of networks were performed as previously described for node degree distribution[5], assortativity[14], and node hierarchical organization[6]. Null-model random networks for correlation profiles of assortativity test were generated by local rewiring algorithm that randomizes a network yet conserves degrees of each node[14,19]. Biological processes were defined by hierarchical clustering of YeastNet described as in[25] or by MCL clustering[26] with the granularity parameter selected so as to balance modularity and proteome coverage. For the GO biological processes network, we connected pairs of GO terms sharing at least one annotated yeast protein to generate a network of 5,587 edges among 1,066 GO biological process terms.

## Acknowledgements

We thank Dr. Petter Holme for sharing the internet dating community dataset, and Dr. Gerald F. Davis for the American company director network dataset. This work was supported by grants from the National Research Foundation of Korea (2010-0017649, 2012M3A9B4028641, 2012M3A9C7050151) to I.L, and from the N.S.F., N.I.H., U. S. Army (58343-MA) and Welch Foundation (F-1515) to E.M.M.

## Author contributions

I.L. and E.M.M. conceived the project. I.L. analyzed data and wrote the manuscript. E.K assisted in implementation of topological analysis algorithms. I.L. and E.M.M. edited the manuscript.

## Additional information

Competing financial interests: The authors declare no competing financial interests.

## Supplementary online material

### Modes of interaction between individuals dominate the topologies of real world networks

Insuk Lee, Eiru Kim, and Edward M. Marcotte

**Supplementary Figure 1.**
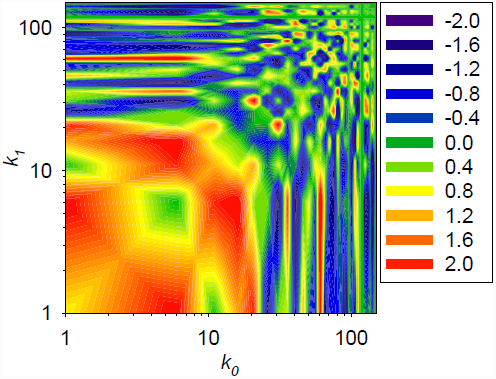
Assortativity analysis for the biological process network, calculated as in Figure 3b.

**Supplementary Figure 2.**
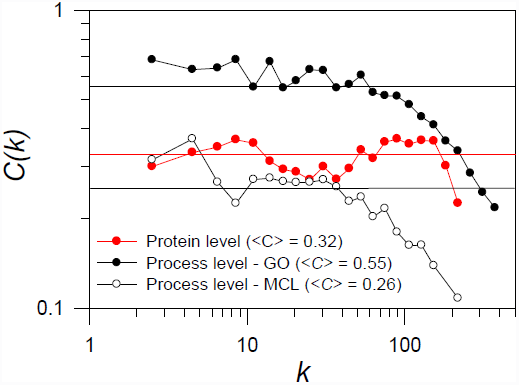
Hierarachy analysis for alternate protein biological process networks. Both a network of previously defined Gene Ontology biological processes (in which edges are defined between 7^th^ level GO-BP annotations based on sharing of annotated proteins; labeled “Process level – GO” in the plot) and the network of biological processes inferred by MCL clustering of a functional protein network (Process level – MCL) show hierarchical organization, while the functional protein network itself (YeastNet core) shows a nonhierarchical organization, in which clustering coefficients for a given degree connectivity (C(k)) trend near the average clustering coefficient for the entire network (<C>).

**Supplementary Table 1.** Summary of the networks analyzed in this study.

| Network | Edge representation | Node representation | # node | # edge | $\langle k \rangle$ |
| --- | --- | --- | --- | --- | --- |
| Protein physical interactions<br>(CCSB-YI1)[1] | Contact-centric | Protein | 1,173 | 1,547 | 2.6 |
| Protein functional interactions<br>(YeastNet core)[2] | Task-centric | Protein | 5,053 | 47,000 | 18.6 |
| Protein processes[2] | Contact-centric | Biological process | 331 | 4,348 | 26.3 |
| Dating[3] | Contact-centric | Person | 19,481 | 58,978 | 6.1 |
| Board of directorships[4] | Task-centric | Person | 1,586 | 11,540 | 14.6 |
| Internet routers[5] | Task-centric | Router | 228,298 | 320,168 | 2.8 |
| Internet domains[5] | Contact-centric | Domain | 16,413 | 31,031 | 3.8 |

